# Convolutional neural network model detected lasting behavioral changes in mouse with kanamycin-induced unilateral inner ear dysfunction

**DOI:** 10.1101/2023.11.04.565603

**Authors:** Masao Noda, Shimada Dias Mari, Chortip Sajjaviriya, Ryota Koshu, Chizu Saito, Makoto Ito, Taka-aki Koshimizu

**Author notes:** Corresponding authors: Masao Noda M.D., Ph.D. E-mali Postal address: 3311-1 Yakushiji, Shimotsuke, Tochigi, 329-0498, Japan ORCID ID: Taka-aki Koshimizu M.D., Ph.D. E-mali Postal address: 3311-1 Yakushiji, Shimotsuke, Tochigi, 329-0498, Japan ORCID ID: 0000-0001-5292-7535. Lead contact: Masao Noda M.D., Ph.D. E-mali.

## Abstract

In acute aminoglycoside ototoxicity to unilateral inner ear, physical abnormalities, such as nystagmus and postural alteration, are relieved within a few days by neural compensation. To examine exploratory behavior over an extended period, freely moving behavior of a mouse after unilateral kanamycin injection was recorded in a home-cage environment. A tail was excluded from deep learning-mediated object detection because of its delayed movement relative to the body. All detection results were confirmed by convolutional neural network classification model. In kanamycin-injected mice, total distance moved in 15 minutes increased at 3 days after surgery. Moreover, the injured mouse turned frequently toward healthy side up to 17 days after surgery. Tail suspension and twist toward healthy side induced fast rotation of trunk around longitudinal axis with dorsal bending after 14 days. Our analysis strategy employing deep learning is useful to evaluate neuronal compensatory process and screen a drug candidate with therapeutic potency.

## 1 INTRODUCTION

Ototoxicity of aminoglycoside antibiotics can cause serious side effects in vestibulocochlear organs and lead to dysfunction of balance and hearing[1, 2]. A challenging feature of the aminoglycoside-induced inner ear dysfunction is unpredictability. In contrast to dose-dependent nephrotoxicity and kidney failure by the aminoglycoside, no apparent association was detected between serum concentrations of the drug and the level of vestibulotoxicity[3, 4, 5]. In a clinical situation, close monitoring of a patient administered with aminoglycoside antibiotics is recommended for early detection of the drug side effect and effective monitoring program is under active discussion[4].

To better understand ototoxicity of aminoglycoside, proper animal model and its analysis method are needed. Initial objective sign of inner ear dysfunction by aminoglycoside can start in one side, and unilateral inner ear dysfunction is more difficult to detect than that of bilateral sever disease, especially in children and elderly[6, 7, 8]. Because aminoglycoside antibiotics damage hair cells in the basal turn of the cochlea, initial hearing loss occur in high-frequency range[9]. Acute hearing loss in one side results in semipermanent failure of sound source localization and sound discrimination[10]. Persistent unilateral hearing loss may cause difficulty in complex conversation in noisy environment for socialization, learning and productive working[11]. Clinically, unilateral hearing loss may be caused by several conditions, such as sudden injury, Meniere’s disease, or early stage of bilateral hearing loss. Either surgical or medical treatment may be used, but if there is no improvement, hearing aids or cochlear implants are some of the few effective options[12, 13, 14, 15, 16].

Although hearing loss by aminoglycoside is lasting disability, previous works in rodents revealed that behavioral phenotypes and signs such as nystagmus due to acute unilateral balance dysfunction last for approximately one week after the injury[17, 18]. Dysfunctions of hearing and balance in peripheral origin are known to undergo central compensation process, because these two functions are controlled by the central integration of inputs from the both sides of the peripheral and central components[17, 10, 11, 19]. However, vestibular compensation is poor in some patients, and a relationship between remaining vestibular function and hearing loss requires better knowledge of the recovery process using proper animal model[20, 21].

In this study, aminoglycoside-induced injury was introduced to unilateral inner ear of mouse and compensatory recovery process was examined in open-field test more than one week after surgery. To localize the mouse in the field and trace its exploratory behavior, deep learning analysis of the object tracing program was developed. The advantage of introducing the deep learning model compare to previous array of photo-beam-based detection is precise and versatile analysis of the mouse movements. However, one of difficulties in employing the object detection by machine learning program in animal experiments is constantly changing background environment, if bedding of the cage floor was included. Because the amount of the bedding and materials condition have an important impact on the animal behavior[22, 23, 24], we developed mouse detection model in the cage with normal bedding same as the home cage environment. Furthermore, a difficulty in evaluation of large number of object detection results was overcome by deep learning-mediated image classification model. Our analysis on the spontaneous exploratory behavior detected lasting change beyond one week after surgery in the mouse with unilateral inner ear injury by aminoglycoside.

## 2 METHODS

### 2.1 Animals and surgical procedure

A total of 40 male C57BL/6J mice at 9 weeks old were obtained from CREA Japan Inc. (Tokyo, Japan). All mice were housed in cages, which were covered by wood-shavings as bedding material under a 12-hour light/dark cycle with food and water. The temperature of the animal room was controlled at 23 ± 1*^◦^*C, and the humidity at 55 ± 10%. The protocols of all animal studies were approved by the Committee on Animal Research at Jichi Medical University in Shimotsuke, Japan (approval no. 20006-01). The experiments were conducted in accordance with Public Health Service Policy on Humane Care and Use of Laboratory Animals. We confirm that details of this study are provided in accordance with the ARRIVE guidelines (https://arriveguidelines.org).

The surgical procedure for creating the deafness model has been reported previously[25, 26, 27]. For anesthesia, a mixture of medetomidine, midazolam, and butorphanol was injected into the intraperitoneal space. We always operated on the left ear, leaving the right ear untreated. After creating a retroauricular incision, the tympanic capsule was drilled and the round window membrane (RWM) was exposed. One microliter of 30% kanamycin solution (dissolved in saline) was then gradually injected into the cochlea through the RWM by using a microsyringe (701N; 26s needle, Hamilton Co., Reno, NV, USA). Thirty three operated mice were recovered for the unilateral inner ear dysfunction model. As the control, 18 mice had 0.9% saline injected into the cochlea instead of kanamycin. We separated entire experiment into two parts (Figure 1). In the first series of the experiment, an experimenter analyzed the mouse behavior in the open-field test and unilateral inner ear dysfunction was confirmed by auditory brainstem response (ABR) 14 days after surgery. At the end of these experiments, histological analysis was performed. In the second series of the experiment, behaviors in the open-field test were analyzed by deep neural network (DNN) model at 3, 6, and 17 days after surgery.

**Figure 1:**
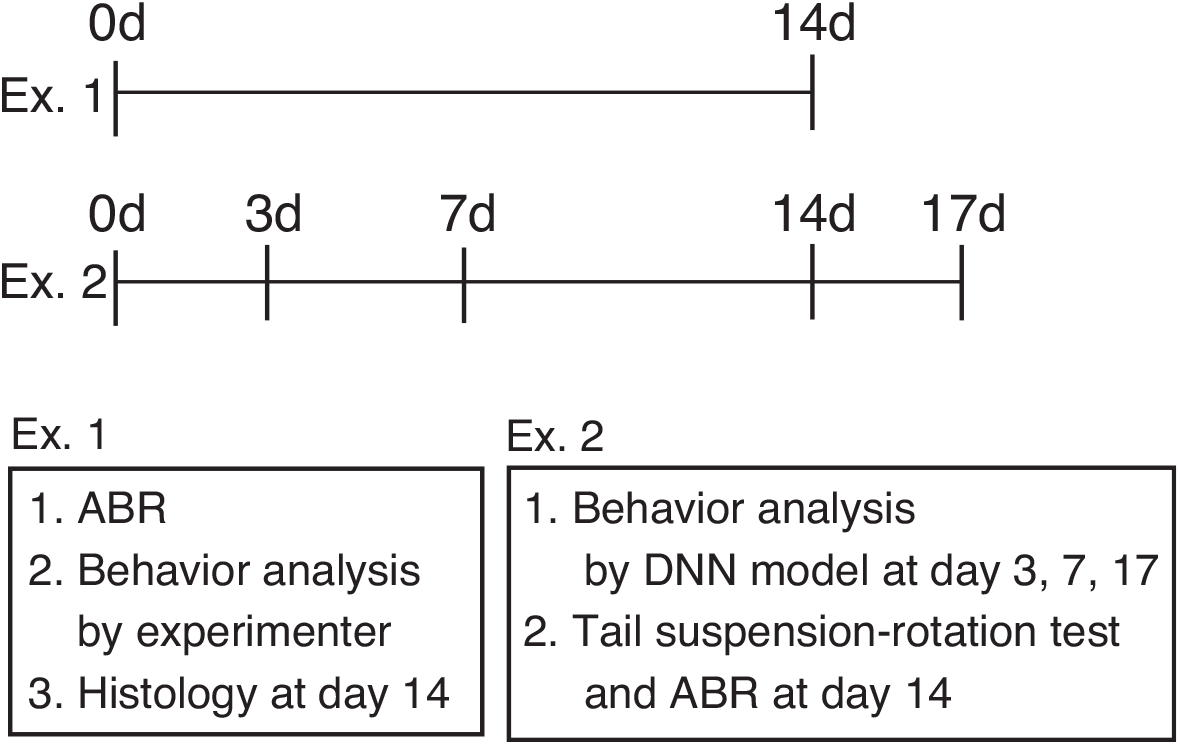
Outline of this study. In the first series of experiment (Ex. 1), unilateral inner ear dysfunction was confirmed by auditory brainstem response (ABR) 14 days after surgery. After the ABR, open-filed test was examined by the experimenter and histological analysis was performed. In the second series of the experiments (Ex. 2), behaviors in the open-field test were analyzed by deep neural network (DNN) model 3, 7, and 17 days after surgery. At 14 days in Ex. 2, tail suspension-rotation test was recorded in video and ABR was measured.

### 2.2 Auditory brainstem response (ABR) measurement

ABR was obtained in a sound-attenuated room 14 days after the kanamycin injection into the cochlea using the procedure as previously reported[28]. To this end, subdermal needle electrodes were placed under anesthesia at the vertex and in the left and right mastoids. The threshold was evaluated by audiometry of click-evoked ABR using Tucker–Davis Technologies and Scope software (PowerLab; AD Instruments Japan, Aichi, Japan).

### 2.3 Histology

After behavioral tests and ABR tests, the mice were perfused with 4% paraformaldehyde in phosphate-buffered saline, and their inner ears were harvested. The temporal bones were locally perfused and fixed in 4% paraformaldehyde overnight at 4*^◦^*C, rinsed in phosphate-buffered saline, and dehydrated for 30 min at each step: once at 30%, 50%, 70%, 85%, and 95%, and twice at 100% ethanol. Clearing was conducted twice, using xylol for 1 h. The specimens were subsequently immersed twice in liquid paraffin (1 h at 42–46*^◦^*C), embedded in a solid paraffin block, and incubated at 46–52*^◦^*C for 1 day. Deparaffinization was performed in a graded ethanol series for 5 min at each step: once at 30%, 50%, 70%, 85%, and 95%, and twice at 100%. The specimens were subsequently rinsed twice with distilled water for 5 min, and cut into 4-µm sections by using the Microm HM340E microtome (Thermo Fisher Scientific, Waltham, MA, USA). The cross-sections were stained with hematoxylin and eosin. Histological analysis and image acquisition were conducted using the Olympus BX51 microscope (Olympus, Kyoto, Japan).

### 2.4 Open-field test

We evaluated the locomotor and exploratory activities of mice in the open-field test in a new polycarbonate cage (18 cm × 26 cm × 12 cm) with heat-treated wood shaving bedding. The behavior was recorded for 15 min by using a video camera (Victor, model GZ-MG77S; JVC, Yokohama, Japan) at 30 fps and at 640 × 480 pixels per image. The behavior at day 14 was evaluated with human observation and at day 3, 7, and 17 with a deep learning model. For observation by human, the number of times of head tossing, sniffing, and touching the wall, and the staying time in the center area were counted three times for each mouse, and the average numbers were used.

### 2.5 Detection of mouse movements

For the automated calculation of mouse movements, sequential images from video records were obtained at five images per second. The locations of mice in the training images were identified and boxed by using LabelImg software (Tzutalin; https://github.com/tzutalin/ labelImg). We used 5682 images with labels, which were randomly partitioned into training/validation (90%), and test (10%) sets. We used transfer learning using single-shot multibox detector_ResNet50 (i.e., SSD_ResNet50), which was pretrained on the COCO 2017 dataset, available from TensorFlow 2 Detection Model Zoo (https://github.com/tensorflow/models/blob/master/research/object_detection/g3doc/tf2_detection_zoo.md)[29, 30]. The training and evaluation steps were undertaken in the TensorFlow version 2.2 environment and TensorFlow Object Detection API (Google, Inc., Mountain View, CA, USA)[31]. For python code, we referred to web site instruction at https://tensorflow-object-detection-api-tutorial.readthedocs.io/en/2.2.0/. The code was developed using a Macbook Pro with cpu support (Apple inc. OS 12.7), and transferred to Ubuntu 20.04 operating system with a Nvidia A6000 GPU, which was established in a conda environment. After transfer learning, our model achieved 87% mean average precision (mAP) value. The XY coordinates of the center of the detected bounding box were the location of the mouse. Distances and angles moved per image were calculated using python and were plotted on a graph using the ggplot2 package in the R statistics environment (version 4.1.2; The R Foundation, Vienna, Austria)[32, 33]. Moved distances during the open-field test were expressed as relative distances, and the maximum width and height of the image were each set as 1.

### 2.6 Evaluation of detection accuracy using deep learning classification model

Success or failure of the mouse detection was evaluated by independently developed deeplearning classification model. To develop the evaluation model, mouse images were subjected to object detection excluding tail and used as images with successful “positive” label for training. Images of unsuccessful detection were manually eliminated. Images of failed detection with “negative” label were generated by randomly placing a bounding box in an image. The size of the bounding box in failed images were fixed at 0.2 x 0.3 of relative hight x width, where hight and width of an image were set as 1. Random placement of a bounding box in an image resulted in inclusion of a mouse image by chance. Such images of unexpected detection were manually eliminated from failed ’negative’ class. The classification model was developed by transfer learning of *Xception* model pre-trained with Imagenet and fine-tuned using custom data of 8000 training, 1600 evaluation, and 1600 test images according to an example code in Tesorflow and Keras website (h**Error! Hyperlink reference not valid.** Validation accuracy and loss values and detection scores were obtained from Tensorbord. Receiver operating characteristic curve (ROC curve) was generated using separately labeled images of a sham mouse 14 days after surgery.

### 2.7 Tail-suspension-rotation test

A mouse was suspended by pinching up its tail, and then the tail was twisted clockwise to rotate mouse trunk. After its rotation was stopped, the tail was twisted counter-clockwise direction to come back to the initial pinch position. The test was recorded in a video as described above and images from the video record was obtained one image per second. The angle between tail and longitudinal axis of the trunk was inspected through the series of images in a test and an image with the smallest angle was selected for the measurement of a degree of the angle by a python program (https://www.higashisalary.com/english/ entry/python-calc-image-angle).

### 2.8 Data and statistical analysis

Data were expressed as the mean ± standard error of the mean (SEM). The R program was used for the Student’s t-tests or Wilcoxon’s tests between the control and kanamycin groups. For the analysis of the time course of open-field tests, time and drug (saline or kanamycin) were set as two factors, and a two-way analysis variance (ANOVA) was calculated. Statistical significance was set at *p <* 0.05.

## 3 RESULTS

### 3.1 Functional and histological confirmation of kanamycin-induced unilateral inner ear injury

The deafness model was confirmed by ABR measurements 14 days after the kanamycin injection into the left cochlea before behavioral analysis. Click-evoked ABR showed a profound lack of the response in the left ear of the kanamycin-injected mouse with the wave threshold reaching to 80–90 dB (Figure 2A). In contrast, the the wave threshold of the saline group was approximately 30 dB (Figure 2A). Also, ABR of the contra lateral side in the kanamycin was within the normal range.

**Figure 2:**
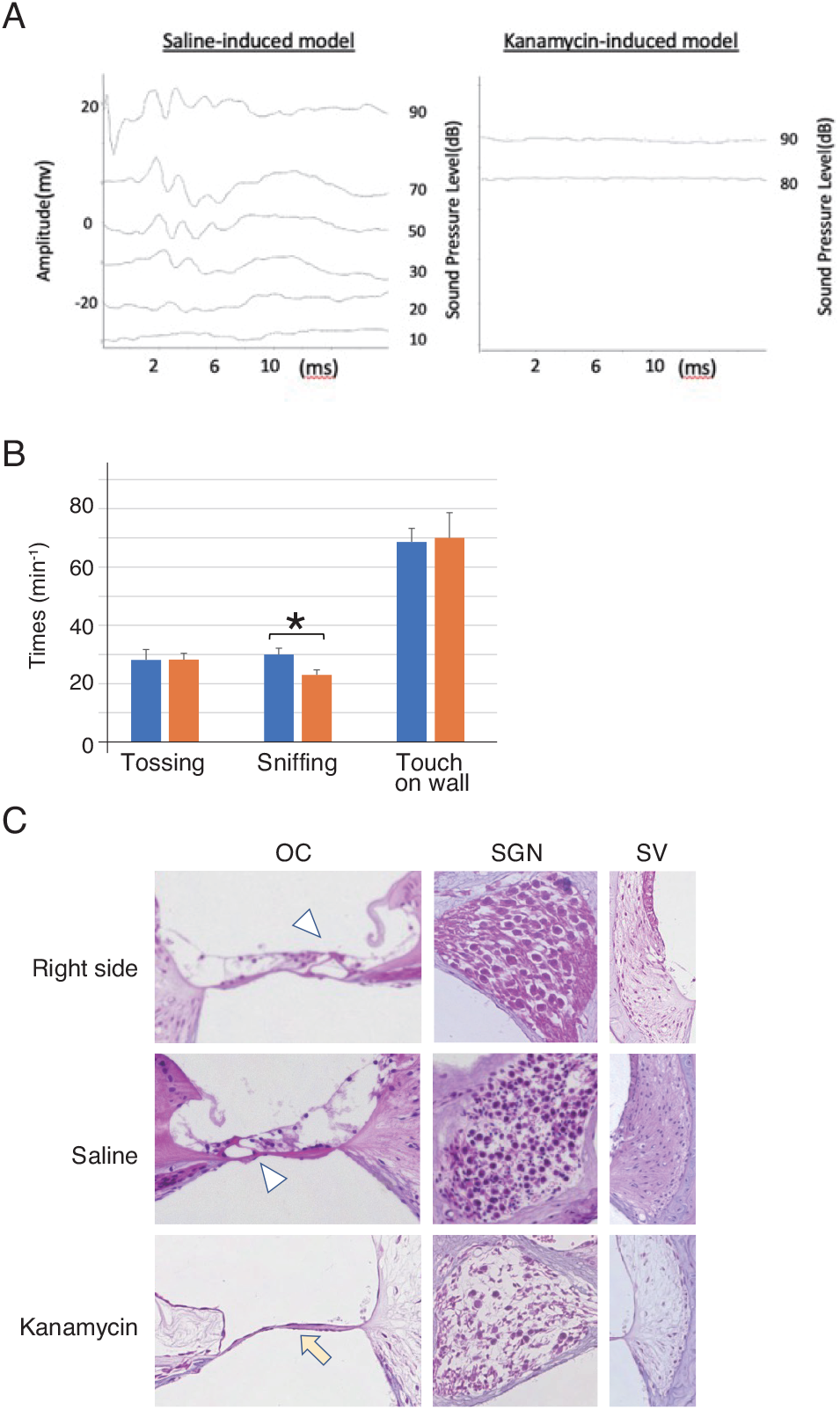
Functional and histological confirmation of the unilateral left inner ear injury. (A) Auditory brainstem response (ABR) was examined in salineand kanamycininjected model mice 14 days after the treatment. The ABR in the saline control mice (left) shows threshold waves around 30 dB. In contrast, in the kanamycin-induced hearing-loss model (right) no waves were detected up to 90 dB. (B) The sniffing behavior was reduced in mice with unilateral inner disorder 14 days after the injection. In the open-field test, activities were counted for 15 min. The bar graphs show the numbers of activities in saline control (blue, n = 9) and kanamycin-injury (orange, n = 7) mice. (C) Cochlear morphology was examined by hematoxylin and eosin staining at 14 days after surgery. Histopathology of cochlear tissue of kanamycin-induced model mice. The images show the cross-section of the cochlea; the organ of Corti (OC), stria vascularis (SV), and spiral ganglion neurons (SGN).

The experimenter analyzed the behaviors of the mice in the video records of the openfield test for 15 min by gross counting. In a 15-min period, there were no significant differences in the numbers of head tosses (28.17 and 28.20 times, *p* = 0.99), touching the wall (68.7 and 70 times, *p* = 0.91), and the counts of entering to the center area (35.6 and 45.5 times, *p* = 0.39) for control and kanamycin-induced model, respectively. However, the count for the sniffing performed by the kanamycin-induced model mice was significantly lower than that by the saline-induced model mice (30 and 23 times for the control and kanamycin-injected models, respectively, *p* = 0.025) (Figure 2B).

The cochlea structure on the right, non-injected side, was intact in both the saline-and kanamycin-injected mice (Figure 2C). In fact, an organ of Corti (OC), a tunnel formed by inner and outer hair cells, the spiral ganglion and stria vascularis were clearly recognized (Figure 2C). In contrast, the cochlear morphology of the left ears of the kanamycininduced injury model showed collapse of the tunnel of hair cells and sever degeneration of the spiral ganglion (Figure 2C).

### 3.2 Establishment of CNN model for the detection of the mouse

After confirming our unilateral inner ear dysfunction model, the mouse behaviors was monitored in open-field test 3, 7, and 17 days after surgery. Exploratory behavior was examined in a clean cage with new wood-shaving bedding. One of the initial difficulty of such analysis in this study was detection of the mouse in changing background image of bedding[35]. In previous open-field experiments, the mouse was detected by computer vision and machine learning method on plane floor with high contract background[36, 37, 38].

By using Tensorflow program for transfer learning, we established the object detection model to localize a mouse in a bounding box and center point in wood-shaving bedding (Figure 3A and 3B). Inclusion of the mouse tail had a significant impact on the locations of the mouse center point and subsequent tracing, because tail was directed to various directions depending on the prior status of the mouse. Inclusion of the tail for mouse detection located the center point relatively posterior position and the center points showed less movements compared to the tracings without the tail (Figure 3C). Furthermore, over all tracing patterns of the control mouse without tail showed clear place preference close to the surrounding walls of the cage (thigmotaxis) (Figure 3D). Therefore, we employed mouse detection method without its tail in the remaining analysis.

**Figure 3:**
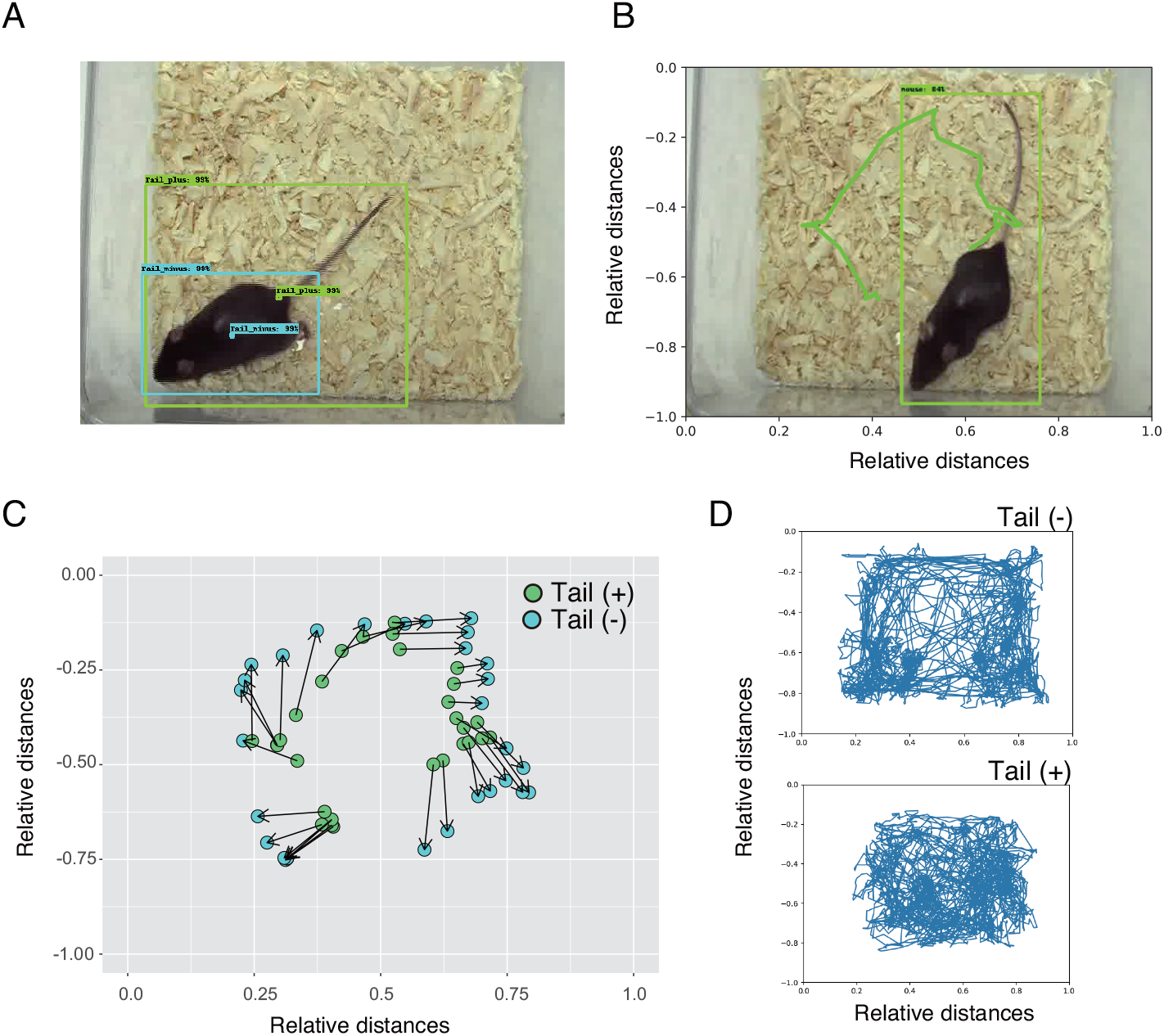
Two types of the detection methods for exploring mouse together with or without its tail in open-field test. (A) Green and blue boxes indicate detection of the mouse with and without its tail, respectively. Center points are indicated in the same colors. (B) An example trace of the mouse movements. In this case, the mouse moved from the position shown in panel (A) to the one shown in (B) in a clockwise direction in 30 images. (C) Differences between the center points with or without tail detection. A series of mouse images from the position of panel (A) to panel (B) were detected by the deep learning model, which include (green) or exclude (blue) the tail. The two types of center points in an image were connected by the arrow. (D) Traces of the mouse movements were different depending on the inclusion status of the tail. From the same video recording of 15 minutes, mouse was detected with or without the tail. Movements of the center point were traced.

### 3.3 Evaluation of object detection efficiency by image classification model

Evaluation of accuracy of object detection results is a difficult task, because a large number of images should be examined. One possible method is employment of deep learning model, which are trained for image classification of the success and failure of bounding box detection. To this end, we performed transfer learning using *Xception* classification model fully trained by Imagenet (Figure 4A). New sets of video recordings of mouse exploratory behavior were prepared as training material. Images from the recordings were subjected to object detection program and images with correct bounding box were labeled as ’positive’. Images of detection failure were removed from training material (Figure 4B *left* panels). To prepare images of failed detection, bounding box was randomly placed in the image. When bounding box with random placement included a mouse by chance, such image was removed from training materials with ’negative’ label (Figure 4B *right* panels). Therefore, an original image of mouse usually produced both ’positive’ and ’negative’ training materials with correct detection and random box placement. During the training and evaluation of the classification model, 4000 and 800 images, respectively, were used per group and obtained high accuracies and low loss values (Figure 4C and 4D). The accuracy and loss values of separated 1600 test images were 0.9994 and 0.0018, respectively. Accuracy of this classification model was examined on 4584 images from an open-field test of 15 min, in which a part of tripod for a video recording was erroneously detected and boxed in123 images (Figure 4F). Classification model correctly classified tripod detection as ’negative’. ROC curve with detection score as a threshold value showed high AUC (area under the curve) value of 0.98 (Figure 4E). Among 232,742 images analyzed in this study, our classification model detected one negative image of actual failure (Figure 4F). Such failed image was excluded from the subsequent analysis. Therefore, our object detection model showed high accuracy according to the deep learning-based classification.

**Figure 4:**
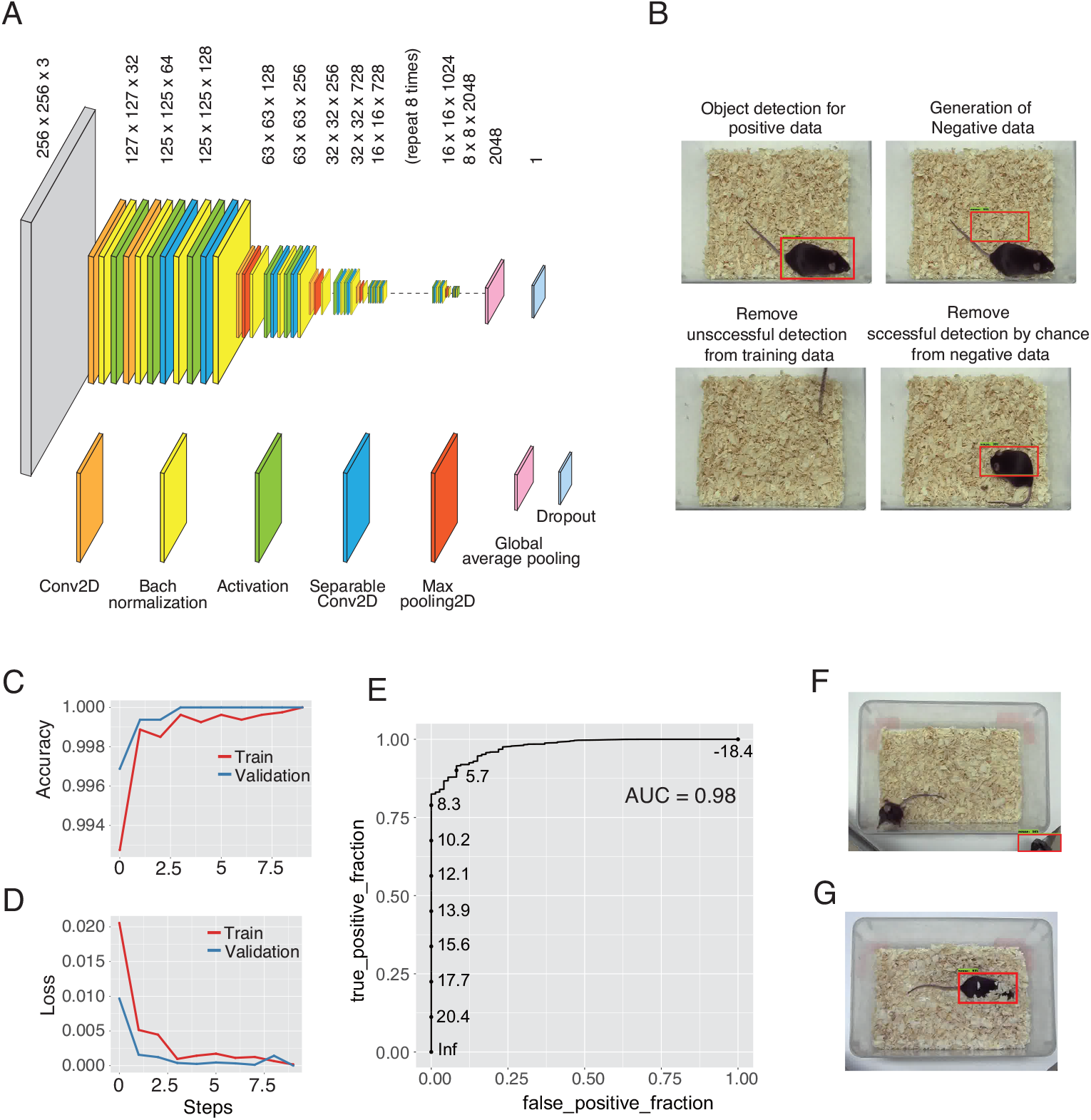
Deep learning-mediated evaluation of object detection results. (A) The *Xception* model was pre-trained on imagenet and the resultant model was used further for transfer learning for image classification to evaluate success/failure of object detection. (B) The model was fine-tuned with custom data set of mouse images with bounding box in the home cage with natural bedding. Images of failed detection were removed from training data. Images with negative detection were generated by placing a box in a random position in an image. If random box was placed on mouse by chance, such image was removed from negative group. (C) and (D) Both positive and negative groups contained 4000 and 800 images for training and validation, respectively. Accuracy and Loss values were shown. (E) Receiver operating characteristic curve (ROC curve) with threshold score was generated from a separate set of open-field experiment for 15 minutes. AUC, area under the curve. (F) One from uniateral inner ear injury and control mice was classified as ’negative’, in which tripod was detected as mouse. This image was excluded from further analysis. (G) The lowest detection score was due to a mouse under the bedding. However, the detection was considered to be successful.

### 3.4 Moved distance was increased 3 days after surgery in kanamycininduced unilateral inner ear dysfunction model

An altered behavioral parameter in kanamycin-treated mice, compared to those of the control mice was found on 3 days after surgery: mice in the kanamycin group explored more distance in the open-field in 15 min than control mice (Figure 5). On the day 3 after surgery, both kanamycin injection and time course were critical determinants on the moved distance: F(85,1) = 84.77, p < 0.0001 for kanamycin/saline injection, and F(85,1) = 88.79, p < 0.0001 for the time course. On the day 7 after surgery, the time course, but not the kanamycin treatment, was a determinant of the moved distances: F(85,1) = 32.81, p < 0.0001 for the time course. In contrast, no statistical difference was detected in moved distances after 17 days after surgery.

**Figure 5:**
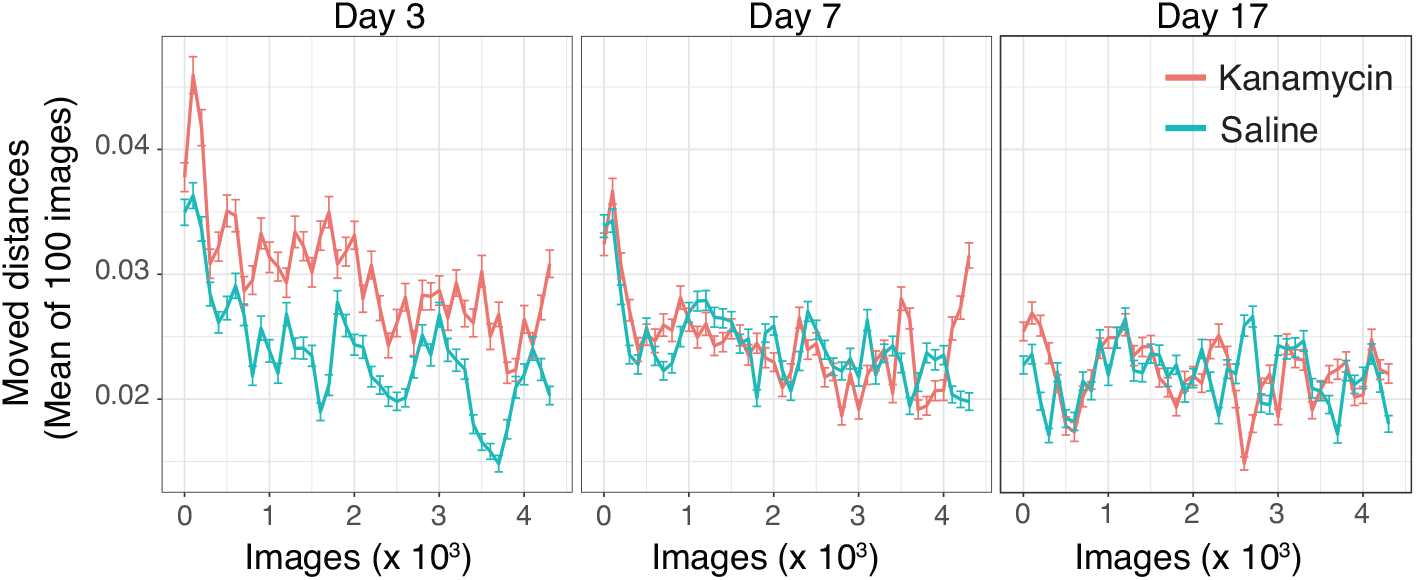
Unilateral inner ear dysfunction model moved longer distances in open-field test 3 days after operation. Video recording of the mouse behavior in the open-field was performed for 15 min 3, 7, and 17 days after operation and the record was converted to images at 5 images/second. Deep learning model localized x and y coordinates of the mouse. Moved distances of the two consecutive images were calculated and averaged distances from every 100 consecutive images were plotted against the sequential numbers of the images. Orange lines show kanamycin injected model (n = 8) and green lines control group (n = 9). Error bars show SEM. The distances of maximum width and hight in an image were set as 1.

### 3.5 Mice in kanamycin-injected group turn to healthy right side more frequently than injured left side

From the video recordings of freely moving mouse in the open-field, we calculated angle of the moving direction (Figure 6A and 6B). Three consecutive images defined an angle.

**Figure 6:**
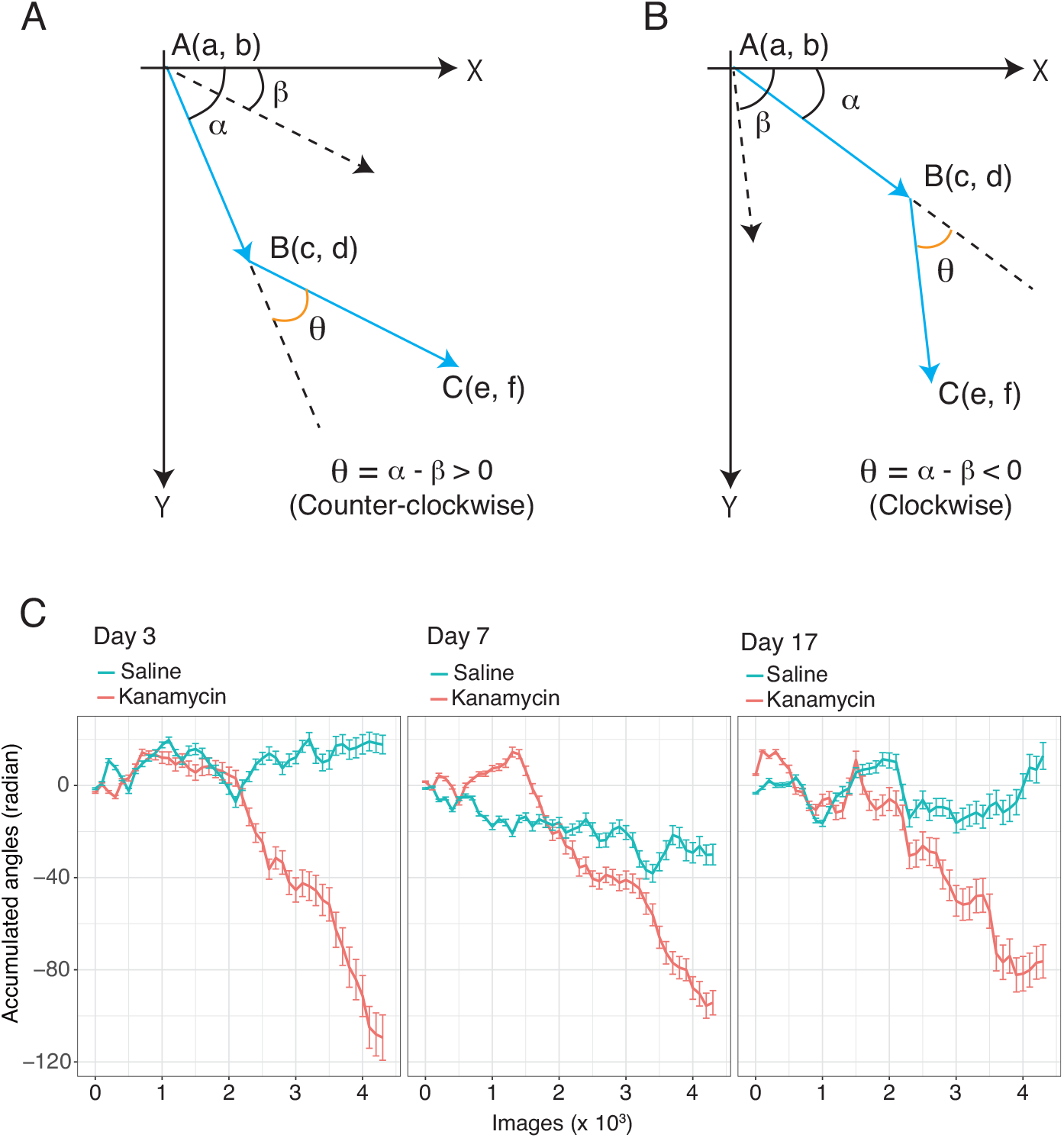
Increase in clockwise rotation of the mouse with kanamycin-induced left inner ear injury. (A) and (B) show calculation procedures of moving direction in a mouse. The mouse proceeded from the point A to B and to C in three consecutive images. The mouse position was localized in an image by object detection program of the deep learning model. The x-y coordinate of the center of the detection box was obtained as the position of the mouse. Angle θ was calculated from angle α minus angle β. The angle α is the angle between x-axis and vector *^−→^* and β is between x-axis and *^−−→^*. From x-y coordinates of the three points of the moving mouse, angles α and β were calculated using atan2 function of R program. Panels A and B also show examples of counter-clockwise, which is an increase in the angle, and clockwise, which is a decrease in the angle, rotations, respectively. The origin of x-y coordinates in the Tensorflow was left upper corner of an image. (C) Accumulated angles during open-field test are shown in the mice with left inner ear injury (orange, n = 8) and in control mice (green, n = 9). The data are the mean ± SEM.

Accumulated angles during the test were close to 0 in control mice, meaning that the mouse turned to right and left equally. In contrast, mouse with left inner ear dysfunction showed clear tendency to turn more frequently to its right hand side, and moved clockwise direction (Figure 6C). Repeated open-field test revealed that the tendency of the mouse with left inner ear injury to turn clockwise was maintained even after 17 days of the surgery (Figure 6C). In the open-field test, both treatment of kanamycin and time course were statistically important determinants of the moving angles (p < 0.001 in day 3, 7, and 17).

### 3.6 Mice with kanamycin moved in clockwise direction more frequently in the cage

The consequence of turning to right more frequently in the left inner ear injury mouse was further examined by visualizing and quantitating mouse movements in the cage with natural bedding. Thigmotaxis is the tendency of the mouse movement close to the walls. We evaluated the direction of the rotation during the thigmotaxis. Lines for the detection of mouse movement direction were set around the middle of the each cage walls in four directions and close to the wall within the 0.25 relative distance. If a mouse crossed any of the four lines in clockwise direction in the next image, the mouse position was colored in green and the count was added plus one (Figure 7). Crossing in opposite, counter-clockwise direction resulted in a purple points and the counts was subtracted one. On the day 7 after surgery, the unilateral injury mouse showed significant increase in the frequencies to turn clockwise direction, which was the side of the healthy ear (Figure 7). The total counts of control and kanamycin mouse were -0.7 ± 6.3 and 27.8 ± 10.5, respectively (*p* < 0.023). Time spent in the center area in the open-field is a measure of anxiety behavior. However, the time in center area was similar levels in control and kanamycin-treated mice.

**Figure 7:**
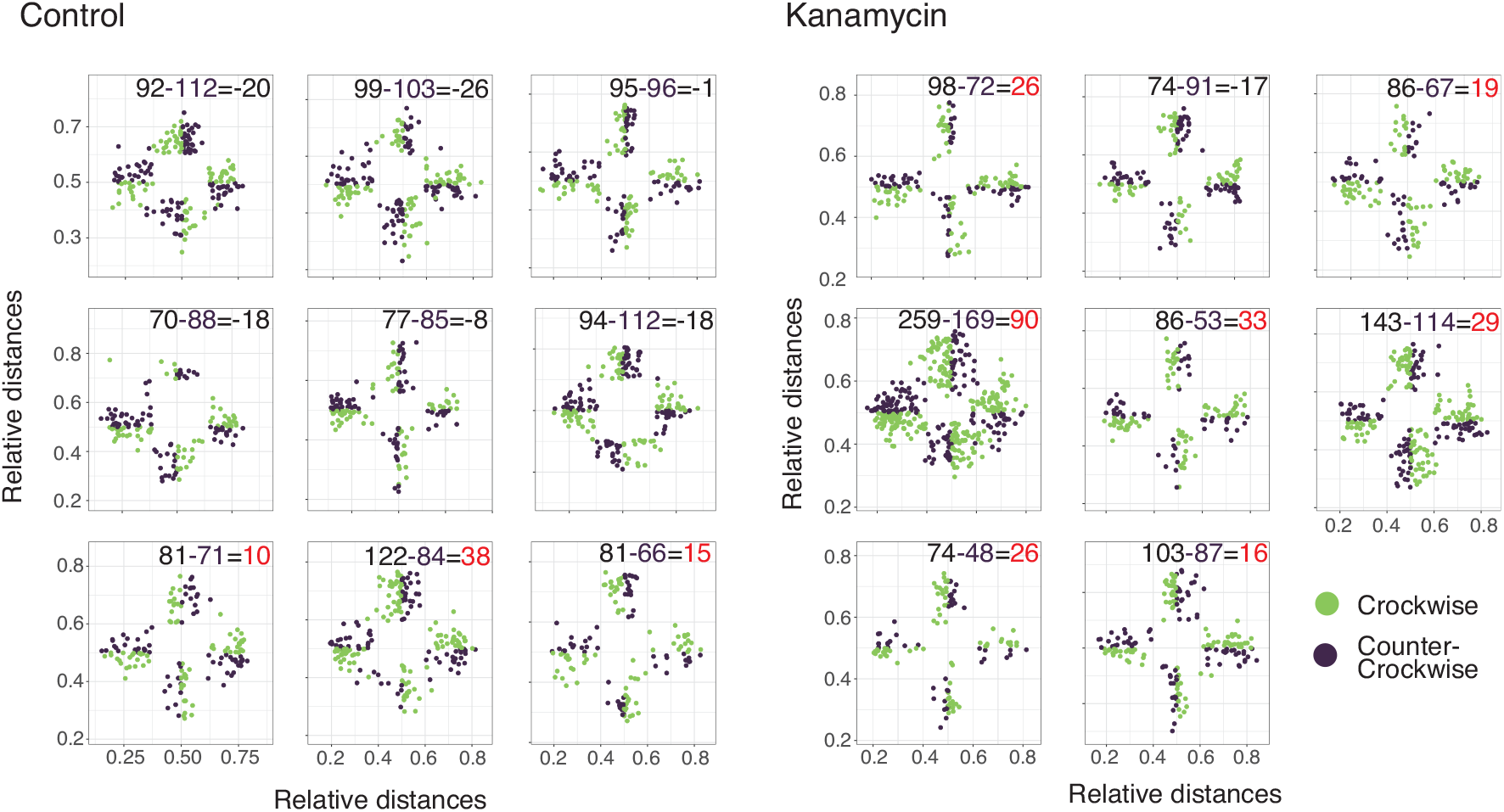
The clockwise movements in the mice with unilateral left inner ear injury. In the open-field images of 7 days post surgery, the position of the mice were recorded as a point if the mice crossed middle line close to cage walls in the next image. Clockwise and counter-clockwise movements were colored in green and purple, respectively. To quantitate the direction of the movement, the crossing in clockwise direction was counted plus one, whereas crossing in counter-clockwise direction was counted minus one. The counts were indicated in the right upper corner area. Red numbers indicate positive total counts and more frequent clockwise crossing of the middle line in the kanamycin group.

### 3.7 Tail suspension-rotation test for the detection of unilateral inner ear injury

In the course of the experiments, we noticed that when an injured mouse was transferred between cages by holding its tail, twisting the tail in clockwise direction resulted in a fast rotation of the body with strong dorsal bend (Supplemental video 1 and 2 for control and injured mouse, respectively). From the video record at 14 days after surgery and measurements of the images from the video, it was found that the degrees of the angles between tail and longitudinal axis of the trunk were significantly smaller in kanamycin-injected mice than those in control mice (Figure 8). Suspending mouse without rotation did not induce dorsal bending or fast rotation.

**Figure 8:**
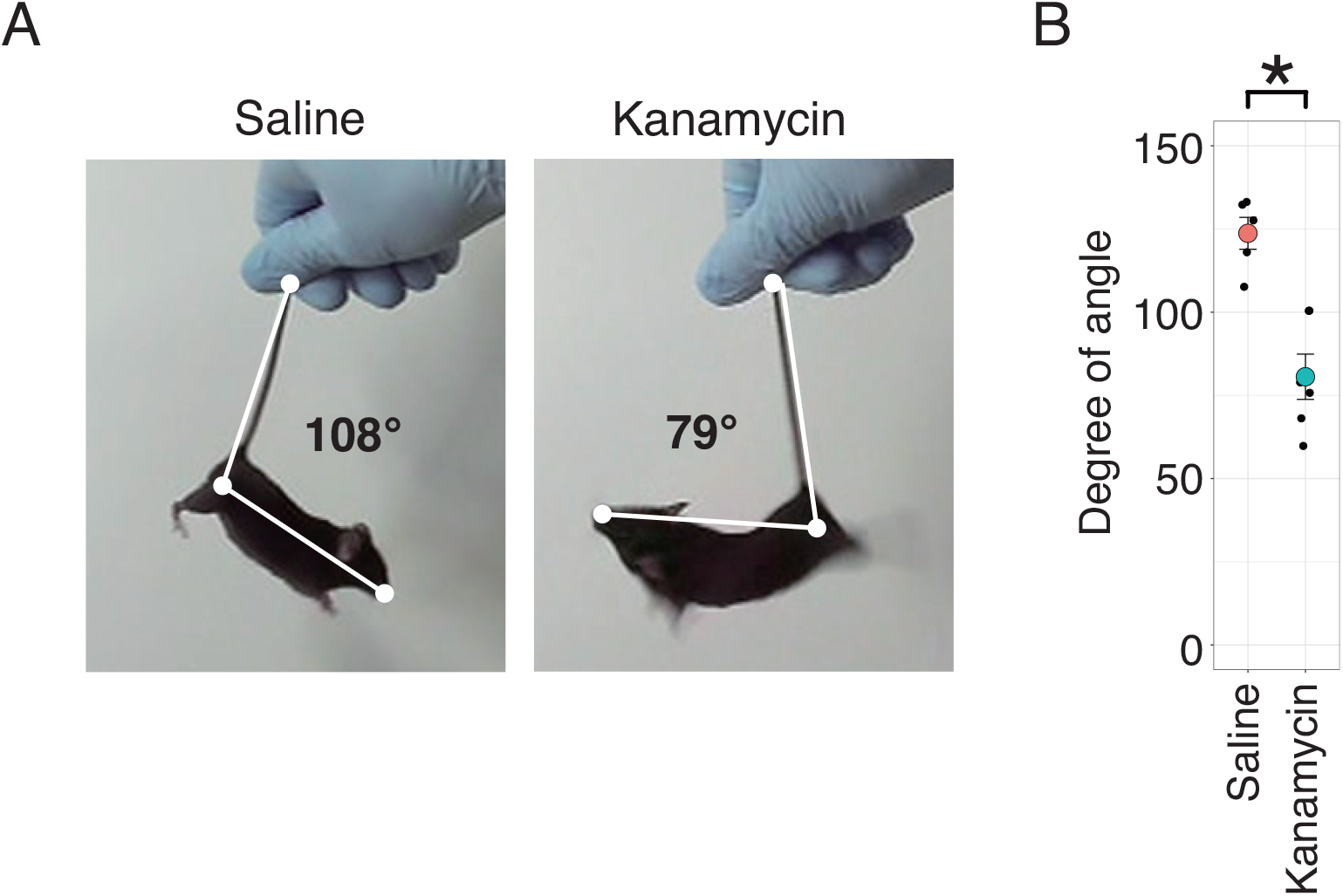
Tail suspension-rotation test for the detection of kanamycin-induced unilateral left inner ear injury. (A) Mice were suspended by their tail and the tail was twisted to rotate the body. Clockwise rotation induced strong dorsal bent in the mouse with left unilateral inner ear injury (right panel), compared to the control mouse (left panel). The degree of the angle between tail and longitudinal body axis was measured from the images of video recordings. (B) The angles in kanamycin-treated mice (n = 6, red) were significantly smaller than those in the control mice (n = 5, green). The test was performed 14 days after surgery. The difference was calculated by two-sample Wilcoxon tests. The error bars show SEM.

## 4 DISCUSSION

In this study, we assessed behavioral abnormalities lasting over one week in mice with unilateral inner ear disorders after kanamycin injection. Unilateral inner ear dysfunction was confirmed by ABR, histology, and positive sign of tail suspension-rotation test. To automate analysis of open-field behavior, efficient CNN model, which enables detection of mouse as an object was developed. For training the model, the mouse tail was excluded from a detection box to obtain center point of the body. This exclusion eliminated influence of tail movement, because inclusion of the tail delayed movement of the mouse. This object detection CNN model was evaluated by image classification CNN model for correct placement of the bounding box. By using evaluated object detection results, our analysis on moved distance in the open-field revealed that kanamycin treatment accelerated travel distance on 3rd day after surgery. More lasting change was clockwise movement in the kanamycin group, which was detected from sequence of images. Also from frequency counts of line crossing, mice with injury in the left ear showed more frequent clockwise movements in the open-field than the movement of opposite direction. In our mouse model, tendency of clockwise movement in kanamycin group lasted beyond one week after surgery. This change in movement direction can be used for a screening purpose of a potential drug that can prevent inner ear injury or facilitate behavioral compensation.

### 4.1 Total distance of movement

Total ambulation distance was increased in unilateral injury mouse only on 3 day post surgery. In previous work by Drago et. al. on young and aged (24 month) male Sprague– Dawley rats, surgical unilateral labyrinthectomy reduced movement score in both age groups of rats[39]. Multiple aspects of experimental conditions were different between our current study and the previous study. To establish unilateral inner ear injury, Drago et. al. used surgical procedure by drilling the left internal ear. Instead, we used round-window approach and kanamycin injection. Movement score in the study by Drago gained the point after both forelegs moved into the next area. However, our measurement was actual distances between detected object in two consecutive images. Moreover, observation period was 15 min in our study instead of two minutes in the study by Drago. In fact, we concluded significant contribution of kanamycin/saline injection from ANOVA by including entire time course and drug treatment for the analysis (Figure 5, day 3).

### 4.2 Frequent movement to the healthy side after unilateral inner ear injury

The behavioral changes lasting for more than one week after injection of unilateral aminoglycoside antibiotics are important findings in this study. When the open-field test was inspected by a human experimenter 14 day after surgery, sniffing activity in the injured mice was less frequent than that in the control mice. Deep leaning model further found that the time course of accumulated angle was markedly different between kanamycin-treated mice and control mice 3, 7, and 17 days after the injury (Figure 6). Left inner ear injury resulted in the accumulation of clockwise turn, which is a healthy side of the right ear, during exploration. Therefore, mouse with unilateral kanamycin injury turned more toward sound source detected in the healthy side of the ear in our study. For calculation of an angle in mouse movements, XY coordinates of mouse positions from three consecutive images were used. In the analysis of movement direction, movements or angles were assumed to have no particular influence on next or future movements or angles. Our results from two directions, calculation of angle and clockwise movement in the cage, indicated the new finding that unilateral inner ear injury caused more frequent movement toward healthy side.

### 4.3 Tail suspension-rotation test

A physical sign was induced in mouse with unilateral inner ear injury by suspending the mouse by the tail and twisting the trail clockwise, which is a direction of healthy ear, resulting in fast clockwise rotation of the trunk. During this rotation, dorsal bending of neck and trunk was observed. In previous study using rat as a model, surgical labyrintectomy or chemical injection of sodium arsanilate into unilateral tympanic cavity induced trunk rotation by lifting rat by the tail[40, 41]. In the previous study on rotation during tail suspension of rat, direction of rotation was in good agreement with the direction of the current study; labyrinthectomy in the right ear induced counter-clockwise rotation by tail suspension[41]. The only one difference between this study on mice and previous ones on rats was that in our study, twisting the tail to clockwise direction was necessary to induce rapid trunk rotation. The rotation sign was sensitive to detect mouse with unilateral inner ear injury; at 14 days after the injury, all 8 mice treated with kanamycin, but none of 9 mice treated with saline, showed this sign.

### 4.4 Lasting change over one week after surgery

These results suggested that unilateral inner ear disorders can cause behavioral abnormalities, even after the acute phase of injury beyond one week. Ito et al. reported that unilateral labyrinthectomy with arsanilic acid causes acute vestibular disorder, which is followed by compensatory mechanisms. They found behavioral abnormalities in terms of circling, dystonia, landing, rotation, and head deviation a few days after surgery, all of which improved by the 7th day when vestibular compensation became active[17].

Binaural hearing helps distinguish between multiple sounds by differences in time, intensity, and frequency spectrum[42, 43]. In unilateral hearing loss, these advantages of binaural hearing are compromised. The significance of these aspects in unilateral hearing loss have been less acknowledged[11]. The unilateral hearing loss in children may have influence on speech perception and communication, language development, and academic performance, and may lead to a potential handicap[44, 45, 46]. Therefore, proper animal model of unilateral hearing loss and its analysis method need to be developed. Lasting disability and time course of the compensation in simple behavior are useful for basic research on the prevention and treatment.

### 4.5 Consistency of object detection CNN model

Our CNN model located center of the object detection bounding box without tail region. This model identified the position and movement of mouse without delay of tail movement. Using this model, moved distance and angle in exploratory behavior were precisely calculated. In previously report on deep learning programs, body parts, but not whole body, are detected as key points, such as paws, snout and eyes, and relative distances between the key points are calculated to extract movement features[47, 35]. However, all key points of body parts are not always detected in images. The paws of a mouse are often covered by beddings in home cage environment. In the recent work using key point detection, lost data points were substituted by related data point[48]. Our analysis model did not fail to place bounding box of 17 mice over 230 thousands images. Therefore, object detection by bounding box has an advantage for behavior analysis in complex environment, such as in the home cage with natural bedding.

Evaluation of object detection results is usually difficult because a large number of images are subjected to the detection. In practice so far, the model is evaluated on a limited number of test images with known object locations, in which detection results were compared with ground truth. We developed deep learning classification model to score and detect success and failure of mouse detection using a new set of images. The classification model was evaluated in ROC curve with high AUC of 0.98. Use of this classification model enabled us to confirm all data of more than 230 thousand images. Therefore, deep leaning classification model trained to detect correct bounding box placement is useful to evaluate a large number of the results of object detection.

### 4.6 Clinical perspective

Detection of unilateral hearing abnormalities from daily behavior in a noninvasive manner is particularly useful when dealing with children and older adults, because they may not be able to identify and accurately describe the hearing problem on their own. Hearing tests for children include other sensory tests, such as ABR and auditory steady-state response (ASSR) tests, which are performed under sedation, and conditioned orientation response audiometry (COR), which observes the behavioral response to a sound33-36. While accuracy is high for objective testing, it must be performed under sedation, and while COR is effective, its accuracy varies greatly depending on the examiner’s ability, the circumstances of the examination, and the type of disease. Thus, training is needed to estimate audiometric thresholds with objective hearing tests35. In addition, although newborns undergo ABR screening soon after birth, there is a high probability that it will be missed, with Johnson et al. reporting that as many as 23% of cases of neonatal hearing loss are missed 37-39. Therefore, the objective evaluations over time are more effective for early detection. Especially, evaluation of behavior by deep learning model may have a possibility to improve detection and treatment effect of unilateral inner ear disorder.

## Acknowledgements

We would like to thank Ms. Yuki Oyama and Ms. Marie Tanaka for their technical assistances.

## Data availability

This manuscript was submitted to bioRxiv. The datasets analyzed in this study can be provided in bioRxiv and Harvard Dataverse[49].

## Author contribution statement

M.N. and T.K. conceived the project and designed the experiments. M.N., S.D.M., S.C., R.K., and T.K. performed the experiments. M.N. and T.K. analyzed data and wrote the manuscript. M.N. and T.K. coordinated and directed the project. All authors reviewed the manuscript.

## Ethics approval statement

Our animal experiments were approved by The Animal Care and Use Committee of the Jichi Medical University.

## References

[1] Dhanireddy, S., Liles, W. C. & Gates, G. A. Vestibular toxic effects induced by once-daily aminoglycoside therapy. Arch Otolaryngol Head Neck Surg 131, 46–8 (2005). URL https://www.ncbi.nlm.nih.gov/pubmed/15655184.

[2] Nakashima, T., Teranishi, M., Hibi, T., Kobayashi, M. & Umemura, M. Vestibular and cochlear toxicity of aminoglycosides–a review. Acta Otolaryngol 120, 904–11 (2000). URL https://www.ncbi.nlm.nih.gov/pubmed/11200584.

[3] Ariano, R. E., Zelenitsky, S. A. & Kassum, D. A. Aminoglycoside-induced vestibular injury: maintaining a sense of balance. Ann Pharmacother 42, 1282–9 (2008). URL https://www.ncbi.nlm.nih.gov/pubmed/18648019.

[4] Konrad-Martin, D., et al. Applying u.s. national guidelines for ototoxicity monitoring in adult patients: perspectives on patient populations, service gaps, barriers and solutions. *Int J Audiol* 57, S3–S18 (2018). URL https://www.ncbi.nlm.nih.gov/pubmed/29157038.

[5] Jiang, M., Karasawa, T. & Steyger, P. S. Aminoglycoside-induced cochleotoxicity: A review. Front Cell Neurosci 11, 308 (2017). URL https://www.ncbi.nlm.nih.gov/pubmed/29062271.

[6] Bowsher, B. Sensorineural deafness following routine transurethral resection of the prostate. BMJ Case Rep 2015 (2015). URL https://www.ncbi.nlm.nih.gov/pubmed/26564118.

[7] Selleck, A. M., Brown, K. D. & Park, L. R. Cochlear implantation for unilateral hearing loss. Otolaryngol Clin North Am 54, 1193–1203 (2021). URL https://www.ncbi.nlm.nih.gov/pubmed/34535281.

[8] Mick, P., Kawachi, I. & Lin, F. R. The association between hearing loss and social isolation in older adults. Otolaryngol Head Neck Surg 150, 378–84 (2014). URL https://www.ncbi.nlm.nih.gov/pubmed/24384545.

[9] Cunningham, L. L. & Tucci, D. L. Hearing loss in adults. N Engl J Med 377, 2465–2473 (2017). URL https://www.ncbi.nlm.nih.gov/pubmed/29262274.

[10] Kumpik, D. P. & King, A. J. A review of the effects of unilateral hearing loss on spatial hearing. Hear Res 372, 17–28 (2019). URL https://www.ncbi.nlm.nih.gov/pubmed/30143248.

[11] Snapp, H. A. & Ausili, S. A. Hearing with one ear: Consequences and treatments for profound unilateral hearing loss. *J Clin Med* 9 (2020). URL https://www.ncbi.nlm.nih.gov/pubmed/32260087.

[12] Nelissen, R. C., Agterberg, M. J., Hol, M. K. & Snik, A. F. Three-year experience with the sophono in children with congenital conductive unilateral hearing loss: tolerability, audiometry, and sound localization compared to a bone-anchored hearing aid. Eur Arch Otorhinolaryngol 273, 3149–56 (2016). URL https://www.ncbi.nlm.nih.gov/pubmed/26924741.

[13] Van de Heyning, P. et al. Incapacitating unilateral tinnitus in single-sided deafness treated by cochlear implantation. Ann Otol Rhinol Laryngol 117, 645–52 (2008). URL https://www.ncbi.nlm.nih.gov/pubmed/18834065.

[14] Blasco, M. A. & Redleaf, M. I. Cochlear implantation in unilateral sudden deafness improves tinnitus and speech comprehension: metaanalysis and systematic review. Otol Neurotol 35, 1426–32 (2014). URL https://www.ncbi.nlm.nih.gov/pubmed/24786540.

[15] Buechner, A. et al. Cochlear implantation in unilateral deaf subjects associated with ipsilateral tinnitus. Otol Neurotol 31, 1381–5 (2010). URL https://www.ncbi.nlm.nih.gov/pubmed/20729788.

[16] Thomas, J. P., Neumann, K., Dazert, S. & Voelter, C. Cochlear implantation in children with congenital single-sided deafness. Otol Neurotol 38, 496–503 (2017). URL https://www.ncbi.nlm.nih.gov/pubmed/28288475.

[17] Ito, T., et al. Vestibular compensation after vestibular dysfunction induced by arsanilic acid in mice. *Brain Sci* 9 (2019). URL https://www.ncbi.nlm.nih.gov/pubmed/31752103.

[18] Wijesinghe, R. & Camp, A. The intrinsic plasticity of medial vestibular nucleus neurons during vestibular compensation-a systematic review and meta-analysis. Syst Rev 9, 145 (2020). URL https://www.ncbi.nlm.nih.gov/pubmed/32552855.

[19] Hamann, K. F. & Lannou, J. Dynamic characteristics of vestibular nuclear neurons responses to vestibular and optokinetic stimulation during vestibular compensation in the rat. Acta Otolaryngol Suppl 445, 1–19 (1987). URL https://www.ncbi.nlm.nih.gov/pubmed/3502519.

[20] McDonnell, M. N. & Hillier, S. L. Vestibular rehabilitation for unilateral peripheral vestibular dysfunction. *Cochrane Database Syst Rev* 1, CD005397 (2015). URL https://www.ncbi.nlm.nih.gov/pubmed/25581507.

[21] Dobie, R. A., Black, F. O., Pezsnecker, S. C. & Stallings, V. L. Hearing loss in patients with vestibulotoxic reactions to gentamicin therapy. Arch Otolaryngol Head Neck Surg 132, 253–7 (2006). URL https://www.ncbi.nlm.nih.gov/pubmed/16549744.

[22] Kawakami, K. et al. Evaluation of bedding and nesting materials for laboratory mice by preference tests. Exp Anim 56, 363–8 (2007). URL https://www.ncbi.nlm.nih.gov/pubmed/18075196.

[23] Freymann, J., Tsai, P. P., Stelzer, H. & Hackbarth, H. The amount of cage bedding preferred by female balb/c and c57bl/6 mice. Lab Anim (NY) 44, 17–22 (2015). URL https://www.ncbi.nlm.nih.gov/pubmed/25526055.

[24] Freymann, J., Tsai, P. P., Stelzer, H. D., Mischke, R. & Hackbarth, H. Impact of bedding volume on physiological and behavioural parameters in laboratory mice. Lab Anim 51, 601–612 (2017). URL https://www.ncbi.nlm.nih.gov/pubmed/29160176.

[25] Heeringa, A. N., Stefanescu, R. A., Raphael, Y. & Shore, S. E. Altered vesicular glutamate transporter distributions in the mouse cochlear nucleus following cochlear insult. Neuroscience 315, 114–24 (2016). URL https://www.ncbi.nlm.nih.gov/pubmed/26705736.

[26] Zeng, C., Nannapaneni, N., Zhou, J., Hughes, L. F. & Shore, S. Cochlear damage changes the distribution of vesicular glutamate transporters associated with auditory and nonauditory inputs to the cochlear nucleus. J Neurosci 29, 4210–7 (2009). URL https://www.ncbi.nlm.nih.gov/pubmed/19339615.

[27] Shimada, M. D. et al. Macrophage depletion attenuates degeneration of spiral ganglion neurons in kanamycin-induced unilateral hearing loss model. Sci Rep 13, 16741 (2023). URL https://www.ncbi.nlm.nih.gov/pubmed/37798459.

[28] Liu, Y. et al. Protection against aminoglycoside-induced ototoxicity by regulated aav vector-mediated gdnf gene transfer into the cochlea. Mol Ther 16, 474–80 (2008). URL https://www.ncbi.nlm.nih.gov/pubmed/18180779.

[29] Liu, W. et al. Ssd: Single shot multibox detector. arXiv e-prints arXiv:1512.02325 (2015). URL https://ui.adsabs.harvard.edu/abs/2015arXiv151202325L.

[30] Lin, T. et al. Microsoft COCO: common objects in context. CoRR abs/1405.0312 (2014). URL http://arxiv.org/abs/1405.0312. 1405.0312.

[31] Huang, J. et al. Speed/accuracy trade-offs for modern convolutional object detectors. In 2017 *IEEE Conference on Computer Vision and Pattern Recognition (CVPR)*, 3296–3297 (IEEE Computer Society, Los Alamitos, CA, USA, 2017). URL https://doi.ieeecomputersociety.org/10.1109/CVPR.2017.351.

[32] Hadley, W. ggplot2: Elegant Graphics for Data Analysis (Springer-Verlag New York, 2016). URL https://ggplot2.tidyverse.org.

[33] R Core Team. *R: A Language and Environment for Statistical Computing*. R Foundation for Statistical Computing, Vienna, Austria (2023). URL https://www.R-project.org/.

[34] Chollet, F. Xception: Deep learning with depthwise separable convolutions. 2017 *IEEE Conference on Computer Vision and Pattern Recognition (CVPR)* 1800–1807 (2016). URL https://api.semanticscholar.org/CorpusID:2375110.

[35] Pereira, T. D. et al. Sleap: A deep learning system for multianimal pose tracking. Nat Methods 19, 486–495 (2022). URL https://www.ncbi.nlm.nih.gov/pubmed/35379947.

[36] Buhler, D. et al. Leptin deficiency-caused behavioral change a comparative analysis using ethovision and deeplabcut. Front Neurosci 17, 1052079 (2023). URL https://www.ncbi.nlm.nih.gov/pubmed/37034162.

[37] Bogachev, M. et al. Video-based marker-free tracking and multi-scale analysis of mouse locomotor activity and behavioral aspects in an open field arena: A perspective approach to the quantification of complex gait disturbances associated with alzheimer’s disease. Front Neuroinform 17, 1101112 (2023). URL https://www.ncbi.nlm.nih.gov/pubmed/36817970.

[38] Sturman, O. et al. Deep learning-based behavioral analysis reaches human accuracy and is capable of outperforming commercial solutions. Neuropsychopharmacology 45, 1942–1952 (2020). URL https://www.ncbi.nlm.nih.gov/pubmed/32711402.

[39] Drago, F., Nardo, L., Rampello, L. & Raffaele, R. Vestibular compensation in aged rats with unilateral labyrinthectomy treated with dopaminergic drugs. Pharmacol Res 33, 135–40 (1996). URL https://www.ncbi.nlm.nih.gov/pubmed/8870029.

[40] Hunt, M. A., Miller, S. W., Nielson, H. C. & Horn, K. M. Intratympanic injection of sodium arsanilate (atoxyl) solution results in postural changes consistent with changes described for labyrinthectomized rats. Behav Neurosci 101, 427–8 (1987). URL https://www.ncbi.nlm.nih.gov/pubmed/3606813.

[41] Hitier, M., Besnard, S., Vignaux, G., Denise, P. & Moreau, S. The ventrolateral surgical approach to labyrinthectomy in rats: anatomical description and clinical consequences. Surg Radiol Anat 32, 835–42 (2010). URL https://www.ncbi.nlm.nih.gov/pubmed/20607261.

[42] Muller, J., Schon, F. & Helms, J. Speech understanding in quiet and noise in bilateral users of the med-el combi 40/40+ cochlear implant system. Ear Hear 23, 198–206 (2002). URL https://www.ncbi.nlm.nih.gov/pubmed/12072612.

[43] Schleich, P., Nopp, P. & D’Haese, P. Head shadow, squelch, and summation effects in bilateral users of the med-el combi 40/40+ cochlear implant. Ear Hear 25, 197–204 (2004). URL https://www.ncbi.nlm.nih.gov/pubmed/15179111.

[44] Steven Colburn, H., Shinn-Cunningham, B., Kidd, J., G. & Durlach, N. The perceptual consequences of binaural hearing. Int J Audiol 45 **Suppl 1**, S34–44 (2006). URL https://www.ncbi.nlm.nih.gov/pubmed/16938773.

[45] Ruscetta, M. N., Arjmand, E. M. & Pratt, S. R. Speech recognition abilities in noise for children with severe-to-profound unilateral hearing impairment. Int J Pediatr Otorhinolaryngol 69, 771–9 (2005). URL https://www.ncbi.nlm.nih.gov/pubmed/15885329.

[46] Noh, H. & Park, Y. G. How close should a student with unilateral hearing loss stay to a teacher in a noisy classroom? Int J Audiol 51, 426–32 (2012). URL https://www.ncbi.nlm.nih.gov/pubmed/22329567.

[47] Mathis, A. et al. Deeplabcut: markerless pose estimation of user-defined body parts with deep learning. Nat Neurosci 21, 1281–1289 (2018). URL https://www.ncbi.nlm.nih.gov/pubmed/30127430.

[48] Porter, J. D., Pellis, S. M. & Meyer, M. E. An open-field activity analysis of labyrinthectomized rats. Physiol Behav 48, 27–30 (1990). URL https://www.ncbi.nlm.nih.gov/pubmed/2236275.

[49] King, G. An introduction to the dataverse network as an infrastructure for data sharing. Sociological Methods and Research 36, 173–199 (2007).

